# QuadSeq: A duplicated-duplex DNA-synthesis technology for high-fidelity nanopore sequencing of linear DNA amplicons

**DOI:** 10.64898/2026.11.17.767292

**Authors:** Marcus Wäneskog, Knud Nor Nielsen, Achille Julienne, Carlos G. Acevedo-Rocha, Emre Özdemir, Emil Damgaard Jensen

**Author notes:** To whom correspondence should be addressed: Marcus Wäneskog, or Emil D. Jensen.

## Abstract

Long-read nanopore sequencing is a highly versatile technology that is currently constrained by high error-rates, which can impair accurate genotyping of complex DNA samples. Common mitigation strategies based on creating consensus sequences from tandem duplications synthesized by rolling-circle or multiple-displacement amplification are prone to create both chimeric DNA molecules and require an initial DNA-circularization. Here we present QuadSeq, a three-step DNA-synthesis technology that can convert linear DNA amplicons into duplicated DNA duplexes that encode two forward and two reverse strand copies of a sequence-of-interest. Intramolecular consensus sequences generated from QuadSeq duplicated DNA duplexes yielded a median single-amplicon accuracy of up to 0.9992 (Q31), without requiring a cumbersome DNA-circularization of the amplicon-of-interest. We can also show that the primary bottleneck for achieving accurate nanopore amplicon sequencing is not the mutation rate of the initial PCR-amplification, but the fidelity of the nanopore sequencing. QuadSeq consensus sequences generated from amplicons synthesized by a low-fidelity DNA polymerase (Taq) had a 6.6-fold lower median error-rate compared to amplicon simplex reads synthesized by a high-fidelity DNA polymerase (Phusion U). Going forward, we predict that the QuadSeq DNA-synthesis technology will enable the use of almost any low-fidelity DNA polymerase for high-fidelity nanopore amplicon sequencing.

## Introduction

Oxford Nanopore Technologies (ONT) have made long-read sequencing both portable and user-friendly. Nanopore sequencing is one of the most versatile sequencing approaches and can be utilized for microbial species-profiling^1–3^, virus and bacterial pathogen detection^4–8^, antimicrobial-resistance-gene typing^9–11^, immunogenetics^12, 13^, genotyping of mutant libraries^14–17^, and for many other applications. Yet, nanopore sequencing is still constrained by relatively high error-rates compared to other competing long-read sequencing platforms, such as PacBio Revio and Vega^1^. The current ONT R10.4.1 flow-cell chemistry has a single-read (simplex) median accuracy of approximately 0.9935 (≈Q22), which can complicate the accurate genotyping of rare variants within high-diversity DNA samples (Fig. 1A)^1, 14, 15, 17^. For low-diversity samples, such as a single bacterial genome, a higher sequencing depth can often be used to generate sufficient coverage to address these errors^18^. However, this approach is not feasible when sequencing complex libraries, or when all sequenced mutant variants are near-identical, i.e. only differ by a few nucleotides^15^. For such DNA samples, unique molecular identifiers (UMIs) have previously been used to facilitate a post-sequencing read clustering and consensus sequence generation^1, 15^. Though, this approach requires that each DNA fragment is copied multiple times after the addition of a UMI-barcode, which decreases the theoretical multiplexing ability and data output from each nanopore flow cell^1, 15^. Moreover, both PCR-errors and nanopore sequencing errors can create mutated and/or erroneous UMI sequences that can be incorrectly paired together, complicating post-sequencing clustering of UMIs^19^. This read pairing also requires a non-trivial amount of computational power, which prevent real-time nanopore sequencing analyses^20^. One strategy to overcome these issues is to instead use ONT duplex sequencing, which can increase single-read accuracy by pairing a forward DNA strand with its complementary reverse strand during nanopore sequencing (Fig. 1B)^21^. Duplex sequencing allows for error-corrections that can enable a median single-read accuracy of 0.999 (>Q30)^21^. However, false duplex sequences, i.e. when two unrelated simplex reads are paired together by the ONT algorithm, can occur^3^. Moreover, the number of duplex-reads generated during a standard ONT sequencing run is typically below 10%, thus limiting high-fidelity nanopore duplex sequencing to just a minority of total reads^21, 22^. To increase the fraction of duplex reads, ONT has previously explored an approach where the forward and reverse DNA strands are physically linked by the ligation of a hairpin adapter (Fig. 1C)^23^. Unfortunately, the presence of a DNA-hairpin stimulates a re-annealing of the DNA duplex *in trans* after a nanopore DNA translocation, which impacts the speed that the complementary strand transitions through the pore, and thus the accuracy of the subsequent basecalling (Fig. 1C)^24, 25^. Thus, when both DNA strands are physically linked, the reverse DNA strand have a noticeably lower sequencing accuracy, which largely nullifies the fidelity gains otherwise obtained by generating a consensus sequence from a physically linked nanopore duplex^24, 25^.

**Figure 1.**
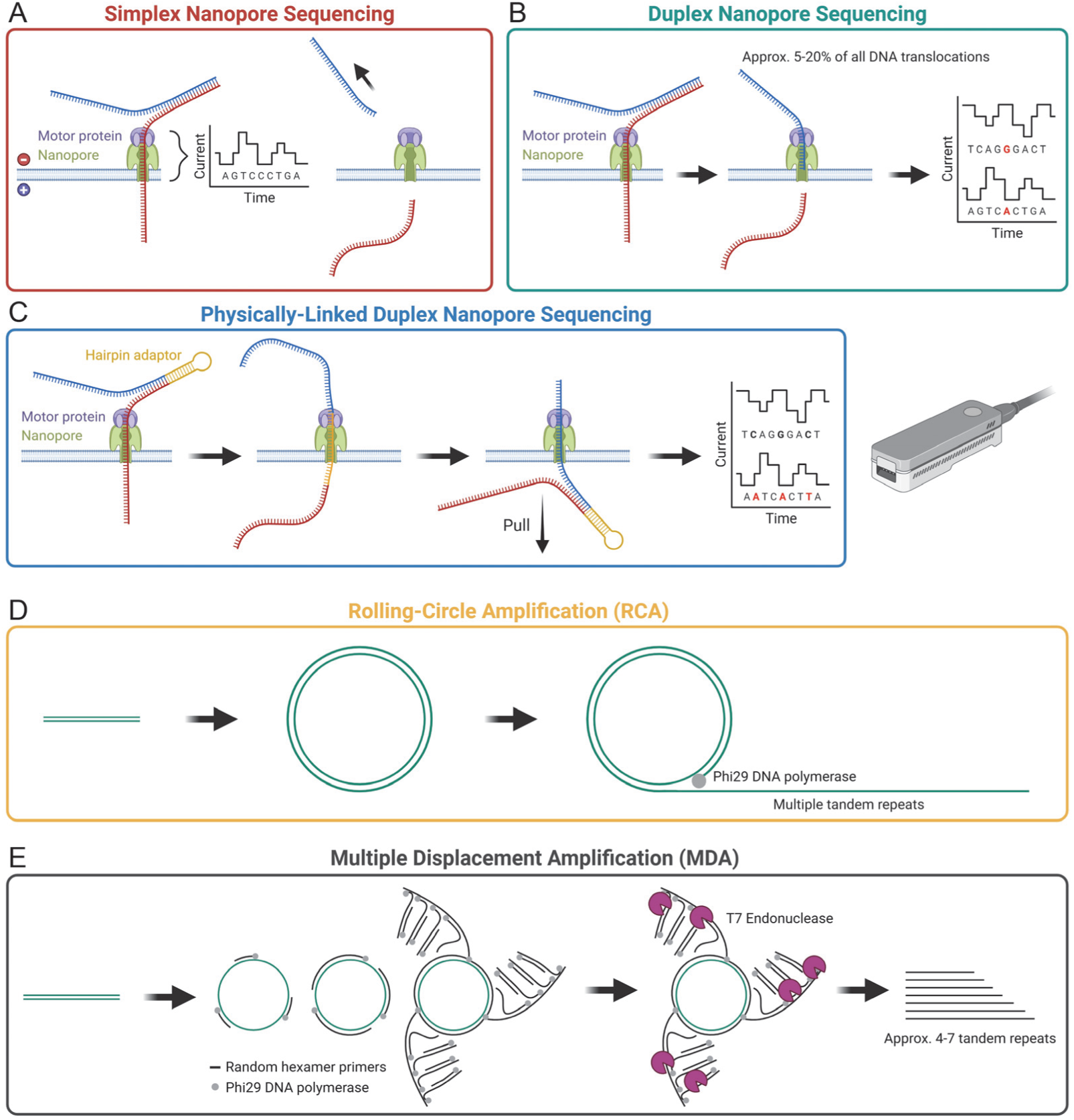
Overview of common strategies used to increase the fidelity of nanopore sequencing. Illustration of conventional ONT **A)** simplex and **B)** duplex sequencing. **C)** Current model for the fidelity-corrupting DNA-re-annealing of inverted-repeats during nanopore sequencing^24, 25^. Illustration of the **D)** RCA and **E)** MDA tandem-duplication DNA-synthesis approaches. After an MDA, the T7 Endonuclease is used to cleave all ongoing DNA-replication junctions to generate linear molecules for sequencing. Illustration created with BioRender.com.

Currently, one of the most common strategies for achieving a higher accuracy during nanopore sequencing, other than relying on coverage, UMIs or duplex sequencing, is by using intramolecular consensus sequencing. During this approach, several tandem duplications of a specific DNA sequence are encoded within a single DNA concatemer, and after nanopore sequencing, all copies are aligned to generate a high-fidelity consensus sequence^26, 27^. This strategy is similar to the circular consensus sequencing (CCS) approach used by Pacific Biosciences (PacBio), which can achieve a single-read median accuracy of 0.9995 (>Q33)^28^. Though, while PacBio CCS utilizes the sequencing information from both the forward and reverse DNA strands to generate a high-fidelity consensus sequence, current nanopore intramolecular consensus sequencing strategies only use the sequence information from either the forward or reverse DNA strand^26–28^. These tandem duplications are synthesized by first circularizing a DNA template and then using either rolling-circle amplification (RCA) or multiple-displacement amplification (MDA) to generate long linear concatemers (Fig. 1D-E)^26, 27^. Yet both of these DNA-synthesis strategies have severe limitations. First, DNA circularization’s can be highly inefficient when using either very short or very long DNA fragments, limiting the possible DNA templates that can be used for generating tandem duplications^29^. Second, multi-branched strand-displacement-mediated DNA amplifications are intrinsically prone to create large DNA rearrangements and chimeric DNA molecules because of both random and sequence-dependent strand-invasions and the synthesis of ssDNA intermediates^30^. For MDA, chimeric molecules and DNA-rearrangements can often be found in up to 78% of all synthesized DNA fragments^31, 32^. For complex DNA samples, such as microbial community barcoding, polyclonal immune repertoires and DNA mutant libraries, these chimeras can be especially problematic because each recombined DNA molecule could be interpreted as a novel variant yet would be nonexistent in the original DNA sample.

A further, and often underestimated, obstacle for amplicon sequencing comes from the PCR-amplification itself. Many of the most valuable amplicons-of-interest can be found in crude DNA samples, such as blood and soil, which also contain many PCR-inhibitors^33^. Most DNA-purification methodologies that remove these impurities introduce a DNA-enrichment bias that can alter the final DNA-composition compared to the original sample^34–37^. For example, most purification methods isolate long dsDNA fragments at higher efficiencies than small dsDNA or ssDNA fragments^34–37^. This can impair quantitative measurements and obscure valuable sequence information from high-value samples. To overcome this problem, valuable amplicons should be PCR-amplified directly from crude and dirty samples. Though, the DNA polymerases (DNAPs) that would be most desirable for this task are often incapable of synthesizing amplicons in crude samples because they also encode exonuclease-domain(s) with weak-to-medium 3’-to-5’ end-resectioning, and/or a 5’-to-3’ flap-endonuclease activity^33, 38, 39^. Normally, this weak-to-medium DNA-degradation activity is balanced by the enzyme’s 5’-to-3’ polymerase activity^33, 38, 39^. However, in the presence of many DNAP-inhibitors, DNA-synthesis becomes impaired, ensuring that the exonuclease domain(s) becomes dominant, which can result in more DNA-degradation than DNA-synthesis. To overcome this problem, most PCR-amplifications of crude and dirty DNA samples are done utilizing DNAP mutants that lack a 3’-to-5’ exonuclease, and/or a 5’-to-3’ flap-endonuclease domain, such as many engineered mutants of the Taq DNAP^33, 40^. For example, Klentaq, OmniTaq and Omni Klentaq, are >10-fold more active in crude and dirty samples, such as blood and soil, compared to wild-type Taq^33, 40^. However, DNAPs without proofreading activity introduces additional mutations during the PCR-amplification^41^. For nanopore sequencing, these mutations create additional errors to the platform’s already relatively high error-rates, which can further impair accurate genotyping of high-value amplicons found in such crude and dirty samples.

Here we present QuadSeq, a three-step DNA-synthesis technology that converts linear DNA amplicons into duplicated DNA duplexes that encode two forward and two reverse strand copies. By creating an intramolecular consensus sequence from each duplicated DNA duplex, QuadSeq can enable a median single-amplicon accuracy of up to 0.9992 (Q31). In addition, QuadSeq does not require an initial DNA-circularization, nor does it rely on a chimeric-prone RCA and/or MDA DNA-amplification. The QuadSeq technology is specifically optimized for synthesizing amplicons using robust, but low-fidelity DNAPs (Taq), yet the intramolecular consensus sequences that are generated from these duplicated-duplex amplicons have a 6.6-fold lower error-rate compared to amplicon simplex reads synthesized by a high-fidelity DNAP (Phusion U). Thus, QuadSeq can enable a high-fidelity nanopore sequencing even when using a low-fidelity DNA polymerase. Going forward, we expect that the QuadSeq technology will also facilitate accurate genotyping and/or rare-variant detection for DNA samples found in complex, crude and dirty samples, a task that is difficult to achieve using other amplicon-synthesis and long-read DNA sequencing methodologies.

## Results

### Motivation and theory

Nanopore sequencing of inverted DNA repeats, i.e. physically-linked DNA duplexes (Fig. 1C), is known to complicate ONT basecalling due to a reannealing of the DNA duplex *in trans*^24, 25^. We hypothesized that a nanopore sequencing of a duplicated duplex, consisting of two forward and two reverse stand copies of a linear DNA amplicon could partially overcome this problem, as duplicated DNA duplexes would consist of two high-fidelity forward strands and two low-fidelity reverse strands. An accurate intramolecular consensus sequence could thus be derived from the two high-fidelity DNA copies, while the two low-fidelity copies could act as conditional tiebreakers for many difficult-to-assess sequencing errors. In addition, because an amplicon-derived duplicated duplex would be both stable and have a predictable and uniform molecular size, we also hypothesized that the synthesis of such a molecule could enable an easy and targeted DNA purification, significantly simplifying most nanopore library preparations workflows. To validate this hypothesis, we designed a straightforward three-step DNA-synthesis technology consisting of a TelN protelomerase-dependent DNA-end-sealing and isothermal strand-displacement-mediated DNA duplication of a DNA amplicon (Fig. 2A).

**Figure 2.**
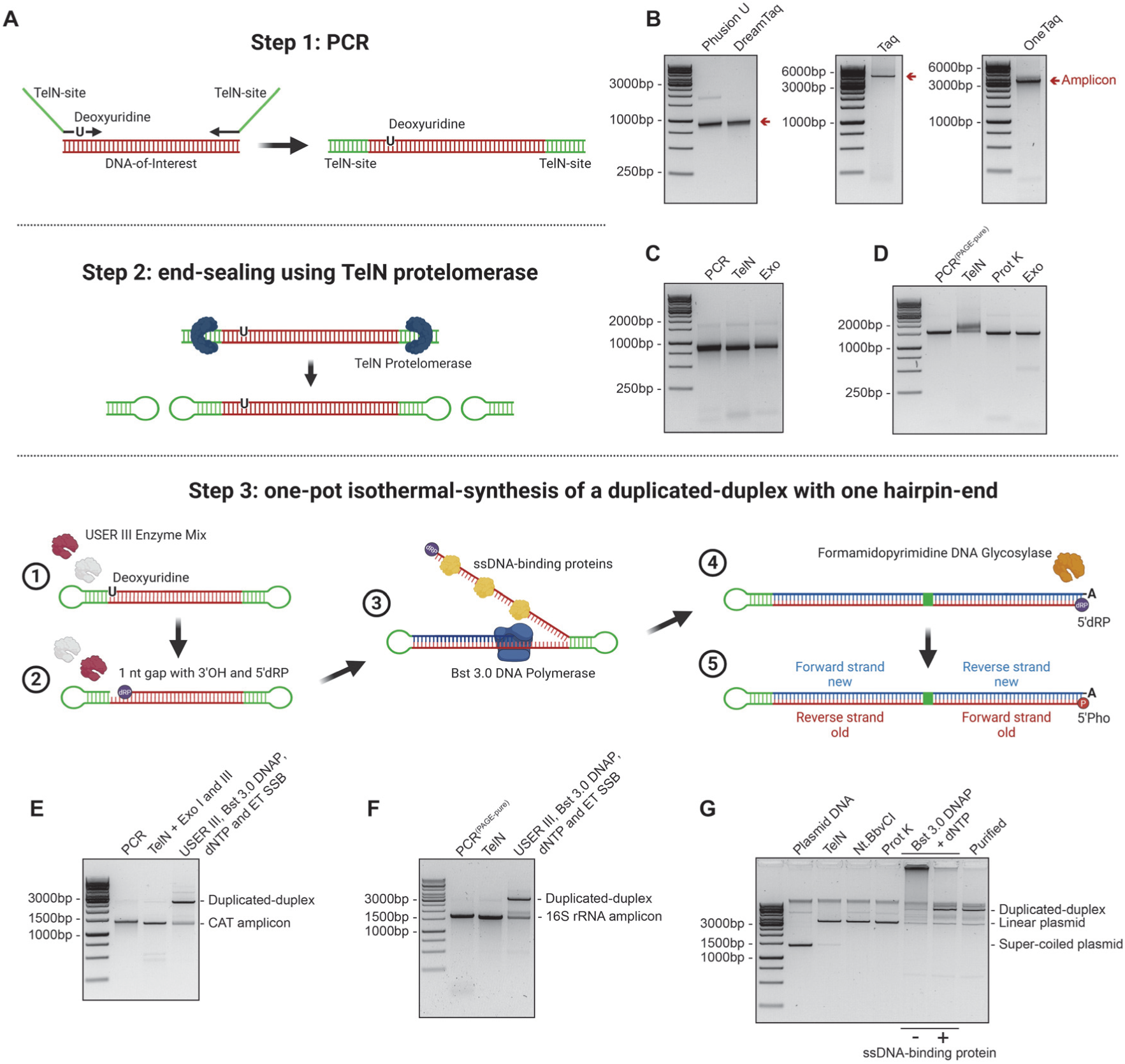
QuadSeq is a three-step DNA-synthesis technology that can convert PCR-modified linear DNA amplicons into duplicated DNA duplexes encoding two forward and two reverse strand copies. **A)** Illustration of the QuadSeq technology. Figure created with BioRender.com. **B)** PCR-amplifications of various DNA regions-of-interest by the Phusion U, DreamTaq, Taq or OneTaq DNAP mixture and QuadSeq-enabled primers containing TelN-sites (both primers) and a deoxyuridine base (only in the forward primer). Evaluation of the efficacy of the TelN end-sealing reaction. Covalently closed amplicons synthesized by **C)** none-purified, or **D)** PAGE-purified QuadSeq primers, were incubated with both Exonuclease I and Exonuclease VIII to assess the fraction of end-sealed molecules. TelN was heat-inactivated before evaluating the efficacy of none-PAGE-purified primers, but enzymatically degraded by a thermolabile proteinase K mutant, before the evaluation of the PAGE-purified QuadSeq primers. A one-pot isothermal reaction can be employed to synthesize a duplicated DNA duplex with one hairpin-end from covalently closed amplicons that were synthesized by **E)** none-PAGE-purified, or **F)** PAGE-purified QuadSeq primers. Purified and covalently closed DNA were incubated with Afu Uracil DNA glycosylase, DNA glycosylase-lyase Endonuclease IV, Bst 3.0 DNAP, extreme thermostable single-stranded DNA binding proteins and dNTPs. Amplicons synthesized by none-PAGE-purified primers were treated with both Exonuclease I and Exonuclease III to degrade none-covalently-closed DNA molecules. **G)** Super-coiled plasmids can be converted to a duplicated DNA duplex if encoding both a TelN-binding-site and a cleavage site for a site-specific nickase. After a TelN treatment, the Nt.BbvCI nickase was used to introduce a DNA-nick downstream of one TelN-hairpin. The resulting linearized and nicked plasmid were incubated with Bst 3.0 DNAP and dNTPs, with or without ssDNA-binding proteins (ET SSB), to synthesis a duplicated plasmid duplex. Thermolabile proteinase K was used to enzymatically degrade both TelN and Nt.BbvCI before the addition of Bst 3.0 DNAP and dNTPs.

### QuadSeq primers are well-suited for both efficient and specific PCR-amplifications

The TelN protelomerase of the *Escherichia coli* bacteriophage N15 binds and cleaves a 56 nt long recognition sequence, thereby creating two covalently closed ends through hairpin formations (Fig 2A)^42^. To exploit this hairpin-forming enzyme for synthesizing a duplicated-duplex, we designed PCR primers targeting different regions-of-interest and incorporated the 56 nt TelN recognition site into the 5’ end of each oligo. The TelN-motif is a semi-perfect inverted-repeat sequence that is highly prone to self-anneal or anneal with other TelN sequences. Thus, we also added 4-5 additional, primer-specific and non-base-pairing nucleotides to the 5’-end of each oligo to prevent a perfect overlap-extension DNA-annealing and primer extension at the 3’-end of all synthesized amplicons during a PCR. In the forward primer sequence, we also introduced a deoxyuridine by replacing a naturally occurring thymine close to the 3’-end, but not directly at the 3’-end of the oligo. We then evaluated whether these deoxyuridine and TelN-site-encoded QuadSeq primers could amplify different sized DNA fragments and whether this primer design was compatible with both high- and low-fidelity DNAPs (Fig. 2B). Both the deoxyuridine-tolerant high-fidelity Phusion U DNAP mutant, and the intrinsically deoxyuridine-tolerant low-fidelity DreamTaq DNAP, could amplify a approx. 900 nt long chloramphenicol acetyl transferase (CAT) gene without any issues (Fig. 2B). The Phusion U DNAP also synthesized a DNA-fusion product, i.e. an overlap-extension CAT amplicon, presumably because the 3’-to-5’ exonuclease domain of the DNAP removed non-base-pairing nucleotides at the primers 3’end(s), which could enable a primer extension even when the two terminal DNA sequences does not perfectly match/anneal^43^. Yet, these chimeric DNA amplicons are inconsequential for successfully synthesizing a QuadSeq duplicated duplex, since each of the DNA chimeras are fused at their terminal TelN-site(s), which are cleaved during the downstream DNA-end-sealing (Fig. 2A). This separates the PCR DNA-fusion product into single amplicons. Moreover, in addition to the Phusion U and DreamTaq DNAPs, we also successfully amplified an approx. 4500 nt long pSEVA231 plasmid backbone, and an approx. 3200 nt long pSEVA3325 plasmid backbone, using deoxyuridine and TelN-site-encoded QuadSeq primers and either the Taq DNAP, or the OneTaq DNAP mixture, which contains both the Taq and Deep Vent DNAPs (Fig. 2B)^44, 45^. Thus, we demonstrate that long TelN-sites can readily be incorporated into PCR primers and that these primers can be successfully utilized by both high- and low-fidelity DNAPs.

### PAGE-purified primers encode the full-length TelN-recognition site

Next, we aimed to verify that TelN-sites incorporated by a PCR amplification were full-length and could facility a TelN-dependent DNA-end-sealing. TelN-encoded amplicons were co-incubated with the TelN protelomerase. After a TelN heat-inactivation, Exonuclease I (ssDNA-specific) and Exonuclease VIII (dsDNA-specific) were added to the reaction to assess the percentage of amplicons with covalently closed and exonuclease-resistant DNA-ends (Fig. 2C). From this experiment, we observed that less than 70% of all amplicons contained closed hairpin-ends. However, because our deoxyuridine and TelN-site-encoded QuadSeq primers were >80 bp in length, we hypothesized that an incomplete oligonucleotides synthesis could be the cause for this partial DNA-end-sealing, as this could result in a high fraction of oligos with incomplete TelN sequences. To test this hypothesis, we designed new deoxyuridine and TelN-site-encoded QuadSeq primers that were polyacrylamide gel electrophoresis (PAGE) purified. DNA amplicons synthesized with these PAGE-purified primers were co-incubated with the TelN protelomerase, followed by a proteolytic TelN degradation using a thermolabile mutant of proteinase K (Fig. 2D). After a proteinase K heat-inactivation, Exonuclease I (ssDNA-specific) and Exonuclease VIII (dsDNA-specific) were added to assess the percentage of amplicons with covalently closed ends (Fig. 2D). This experiment showed that approximately 95% of all amplicons synthesized with PAGE-purified primers had exonuclease-resistant hairpin-ends (Fig. 2D). Proving that QuadSeq-enabled and covalently-closed DNA molecules can efficiently be created directly from amplicons synthesized with PAGE-purified primers, as well as from amplicons synthesized with non-PAGE-purified primers, if an additional exonuclease clean-up step is also performed (Fig. 2C-D).

### Isothermal-synthesis of a duplicated DNA duplex from linear DNA amplicons

Purified and covalently-closed amplicons of a approx. 1200 nt long region encoding a T7 RNA polymerase promotor and CAT gene were converted into a duplicated-duplex using a isothermal and multi-enzyme reaction comprised of Afu Uracil DNA glycosylase (UDG), DNA glycosylase-lyase Endonuclease IV (Endo IV), Bst 3.0 DNAP (Bst 3.0) and extreme thermostable single-stranded DNA binding proteins (ET SSB) (Fig. 2A, E-F). During this one-pot isothermal reaction, the singular deoxyuridine base located in each covalently closed amplicon is excised by the USER III enzyme mixture, which contains both the uracil-base-excision, UDG, and the abasic-site-cleaving, Endo IV enzymes, creating a 1 nt gap with a 5’-deoxyribose phosphate (dRP) and a 3’-hydroxyl DNA-end. Next, the strand-displacement-capable Bst 3.0 DNAP converts each nicked amplicon into a duplicated DNA duplex by extending the DNA-strand from the USER III-created 3’-hydroxyl DNA-end through the downstream TelN-created DNA-hairpin and inverted-repeat sequence, simultaneously as the ET SSB proteins assist in resolving most inhibitory DNA secondary structures (Fig. 2A, E-F). Finally, formamidopyrimidine DNA glycosylase (Fpg) is added to the reaction to convert the USER III-created 5’-deoxyribose phosphate (dRP) into a 5’-phosphate (Pho), which enables an efficient and straight-forward ONT dT-sequencing-adapter ligation.

By using this three-step QuadSeq methodology, we successfully synthesized duplicated DNA duplexes from linear DNA fragments (Fig. 2E-F). We could also observe that duplicated-duplexes could be successfully created from DNA fragments that were amplified either from homogenous DNA samples, such as the CAT gene, but also from heterogenous DNA samples, such as full-length 16S rRNA genes (Fig. 2E-F). A key advantage of the QuadSeq approach is that because each synthesized duplicated duplex is both stable (dsDNA molecule) and has a predictable molecular size (double the size of the original amplicon), the third and final step in the QuadSeq workflow does not need to have a 100% conversion rate (Fig. 2A). All successfully synthesized duplicated duplexes can easily be purified by gel electrophoresis based on their expected molecule size (Fig. 2E-F). This feature of the QuadSeq methodology is in stark contrast to both the RCA- and MDA-dependent tandem DNA duplication-synthesis strategies, which both create a heterogeneous mixture of ssDNA and dsDNA molecules with a substantial diversity in sizes, complicating any targeted DNA purification^26, 27^. In addition, because the duplicated-duplex DNA-synthesis occurs in a one-pot isothermal reaction, it is also possible to titrate the USER III enzymes to stagger the DNA-synthesis. This can minimize the creation of chimeric DNA molecules, because only a small fraction of the QuadSeq amplicons are actively replicated at any given time.

### Isothermal-synthesis of a duplicated plasmid duplex

The QuadSeq methodology was specifically developed to synthesize duplicated DNA duplexes from amplicons, as this enabled a staggered DNA-synthesis and the insertion of a sequence-independent DNA-nick-site (deoxyuridine) in each amplicon via the forward primer. Similarly, we reasoned that this methodology could also be utilized to achieve a synchronized duplicated DNA duplex synthesis, using an enzymatic and sequence-dependent DNA-nicking strategy, which could be used to convert a plasmid template into a duplicated plasmid duplex (Fig. 2G). To test this hypothesis, we utilized a approx. 3200 nt long pSEVA3325 vector encoding both a TelN-recognition site and a Nt.BbvCI/Nb.BbvCI nickase cleavage site. The TelN protelomerase was co-incubated with this plasmid, and after a TelN heat-inactivation, Nt.BbvCI was used to introduce a DNA-nick downstream of one of the TelN-hairpins (Fig. 2G). The Nt.BbvCI nickase, and the previously heat-inactivated TelN enzyme, were then both proteolytically degraded, using a thermolabile mutant of proteinase K. Following a proteinase K heat-inactivation, the resulting linearized and nicked plasmids were co-incubated with Bst 3.0 DNAP and dNTPs, with or without ssDNA-binding proteins (ET SSB), to synthesis a duplicated-duplex (Fig. 2G). This experiment showed that replacing the sequence-independent deoxyuridine cleavage site with a sequence-dependent nickase cleavage site (Nt.BbvCI) could also enable a duplicated DNA duplex synthesis of a plasmid containing both a TelN- and nickase-cleavage-sites (Fig. 2G). However, this alternative approach requires that the DNA sequence of interest does not already encode the cleavage site of the chosen nickase. We could also confirm that ssDNA-binding proteins are essential for successfully synthesizing a duplicated-duplex if the DNA template is either large, contains complicated DNA secondary structure, or if all DNA molecules are replicated simultaneously during the QuadSeq reaction (Fig. 2G).

### Nanopore sequencing of a duplicated DNA duplex

Unlike the long-read PacBio platforms, Revio and Vega, which sequence DNA by synthesis, ONT reads a DNA molecule while it transits a nanopore as a single-stranded chain. This unfortunately means that nanopore sequencing can be highly sensitive to the formation of complex DNA secondary structure *in trans*, since this can affect the speed at which the DNA translocates through the nanopore (Fig. 1C)^24, 25^. However, a QuadSeq-synthesized duplicated DNA duplex can partially overcome this inhibitory re-annealing by virtue of its unique DNA structure, i.e. the forward-reverse-forward-reverse DNA arrangement. We observed that when the first forward DNA strand in the duplicated duplex transited the nanopore, it was read with the same relative accuracy as a normal simplex read (Fig. 3A-B). When the first reverse DNA strand then transited the nanopore, two theoretical inverted-repeat DNA re-annealing events are possible, one *in cis*, and one *in trans* (Fig. 3A). Yet, presumably because of the DNA helicase activity of the nanopore motor protein, we only observed that the inverted-repeat re-annealing *in trans* affected the DNA translocation speed (i.e. the basecalling accuracy) of the first reverse DNA strand (Fig. 3B). Thus, supporting previous reports on nanopore sequencing of physically linked DNA-duplexes (Fig. 1C)^24, 25^. Nevertheless, by exploiting the unique Forward-Reverse-Forward-Reverse structure of the QuadSeq molecule, it is possible to overcome this inverted-repeat DNA re-annealing, because after the first reverse DNA strand has translocated through the nanopore, the inverted-repeat sequence becomes fully formed and the molecular “pulling” effect stops (Fig. 3A-B). When the second forward DNA strand then transitions through the nanopore, it can be read without any constraints, allowing for high-fidelity basecalling (Fig. 3B). Finally, when the second reverse DNA strand transitions through the nanopore, is also becomes corrupted due to a similar inverted-repeat DNA re-annealing event *in trans* (Fig. 3A-B). Thus, nanopore sequencing of QuadSeq duplicated DNA duplexes can generate two high-fidelity copies of the forward DNA strands and two low-fidelity copies of the complimentary/reverse DNA strands (Fig. 3B).

**Figure 3.**
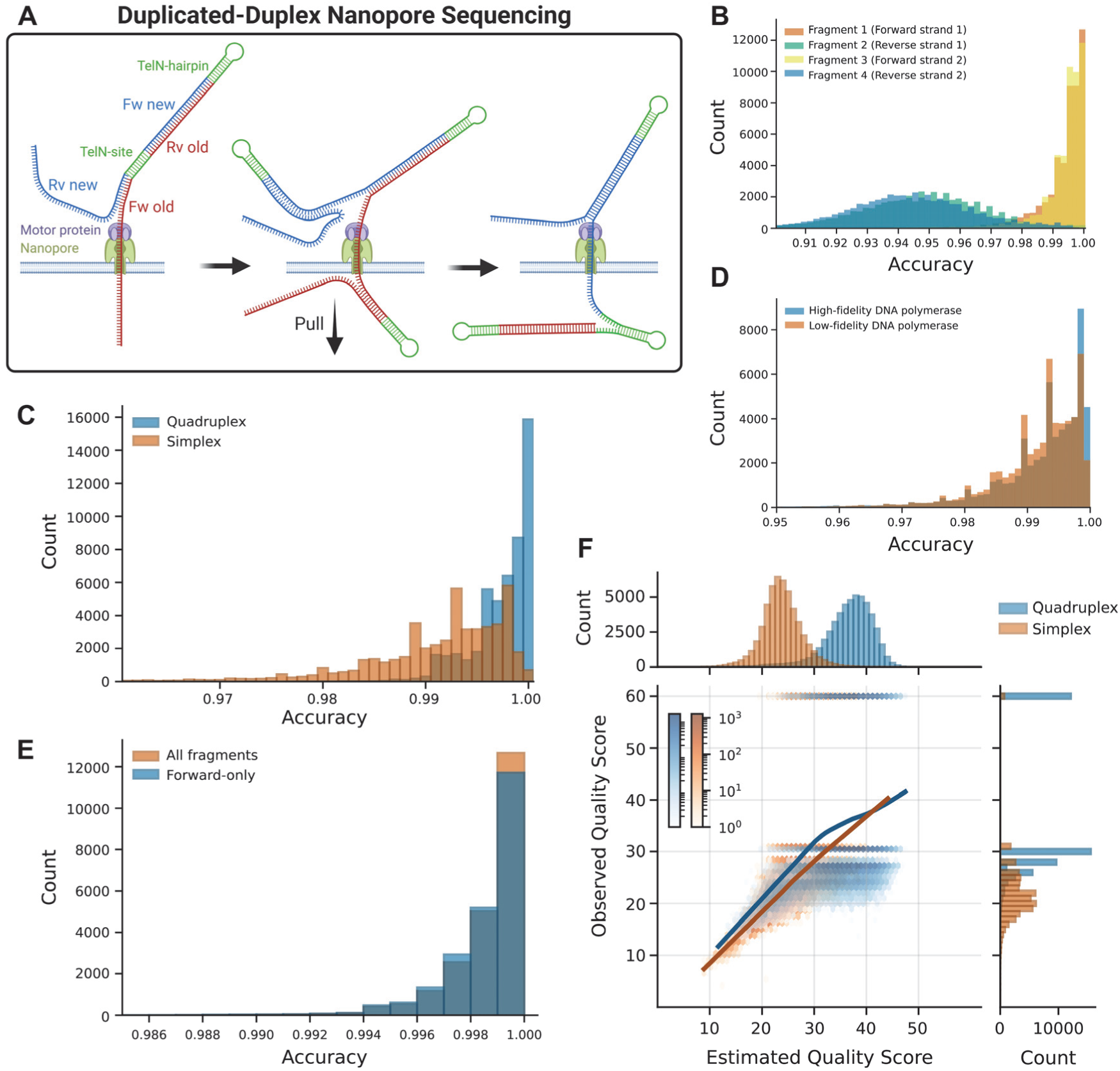
Nanopore sequencing of QuadSeq duplicated DNA duplexes increase single-amplicon accuracy by partially overcoming the fidelity-corrupting DNA-re-annealing of inverted-repeats *in trans*. **A)** Illustration of our proposed model for duplicated-duplex nanopore sequencing. Figure created with BioRender.com. **B)** The basecalled forward DNA strands in each duplicated duplex have a higher median accuracy than both complementary/reverse DNA strands. **C)** Intramolecular consensus sequences generated from QuadSeq duplicated DNA duplexes (Quadruplex) can increase the fraction of error-free single-amplicons by >9-fold. **D)** Simplex nanopore sequencing of amplicons synthesized by either the low-fidelity Taq DNAP, or the high-fidelity Phusion U DNAP, show similar accuracies. Though, the fraction of error-free amplicons is higher when the high-fidelity Phusion U DNAP is used to amplify the region-of-interest. **E)** Filtered full-length QuadSeq intramolecular consensus sequences can reach a median single-amplicon accuracy of 0.9992 (Q31), and the complementary DNA strands can provide conditional error corrections. **F)** Average estimated Phred scores, derived directly from basecalled ONT FASTQ reads, shows a strong correlation with the observed Phred scores of each QuadSeq consensus sequence. A Phred score of 60 denotes error-free amplicons.

### Intramolecular consensus generation of nanopore-sequenced duplicated DNA duplexes

We theorized that both forward DNA strands basecalled after a QuadSeq duplicated-duplex nanopore sequencing could be used to generate a high-fidelity consensus sequence, while the two error-prone reverse DNA strands could potentially be used as conditional tiebreakers for many difficult-to-assess sequencing errors. To test this hypothesis, we created a Nextflow pipeline (DSL2) that generated amplicon consensus sequences from ONT-basecalled FASTQ reads of each duplicated DNA duplex. This consensus generation used a log-likelihood approach to select the base, or gap, with the highest quality-weighted log-likelihood based on the ONT Phred quality scores, assigning the mean score of the neighboring bases to gaps. Using this pipeline, we generated nanopore QuadSeq consensus sequences for duplicated DNA duplexes that were created from a approx. 1200 nt long DNA amplicon (Fig. 2E). We then evaluated the accuracy of these QuadSeq consensus sequences in relation to nanopore simplex reads of the same amplicon synthesized by a conventional PCR (Fig. 3C). QuadSeq duplicated DNA duplexes amplified using the low-fidelity Taq DNAP generated consensus sequences with a median accuracy of 0.9975, corresponding to 2.5 errors per 1000 nt. While Taq-synthesized simplex amplicons had a median accuracy of 0.9930, corresponding to approximately 7 errors per 1000 nt (Fig. 3C, Table 1). The fraction of single amplicons with an accuracy greater than 0.999 (Q30) was 0.0492 for simplex reads (Taq DNAP) and 0.3221 for QuadSeq consensus sequences (Table 1). Almost 13% of all QuadSeq reads were error-free, compared to only 1.4% of simplex reads (Taq DNAP). This quantification showed that QuadSeq duplicated-duplexes can decrease the error-rate of single-amplicons by almost 3-fold, the fraction of high-fidelity reads (>Q30) by more than 6-fold, and the fraction of error-free amplicons (1200 nt) by more than 9-fold, compared to nanopore simplex reads (Table 1). Yet these QuadSeq-synthesized duplicated DNA duplexes were created using DNA that was amplified with a low-fidelity DNAP (Taq). Thus, we next sought to investigate the relative number of mutations that the Taq DNAP introduced during a PCR, compared to the high-fidelity Phusion U DNAP, and how this would impact the perceived nanopore sequencing accuracy. The same approx. 1200 nt long CAT amplicon was PCR-amplified using the Phusion U DNAP and then sequenced by ONT (Fig. 3D). From this analysis, we did not observe that the low-fidelity Taq DNAP introduced a high abundance of PCR errors, which would be interpreted as nanopore sequencing errors when aligned to a reference sequence. The median accuracy of Phusion U-synthesized amplicons (high-fidelity) was 0.9940, corresponding to approximately 6 errors per 1000 nt (Fig. 3D, Table 1). While the fraction of reads with an accuracy greater than 0.999 (Q30) was 0.1273, and the fraction of error-free reads were 0.0513 (Table 1). Thus, we observed that the error rate of ONT sequencing were much higher than the mutation rate of either the Taq or Phusion U DNAP, and QuadSeq consensus sequences generated from duplicated DNA duplexes synthesized by the low-fidelity Taq DNAP had a median error-rate that was 2.4-fold lower than nanopore simplex reads synthesized by the high-fidelity Phusion U DNAP (Table 1). The fraction of high-fidelity (>Q30) QuadSeq reads was also 2.5-fold higher, and the fraction of error-free reads were 2.4-fold higher (Table 1). This data clearly demonstrates that the QuadSeq technology can increase nanopore sequencing fidelity sufficiently to overcome the higher relative mutation rate of the Taq DNAP and also proves that the primary bottleneck for nanopore amplicon sequencing is not the mutation rate of the PCR-amplification, but the accuracy of the nanopore sequencing reaction.

**Table 1.**
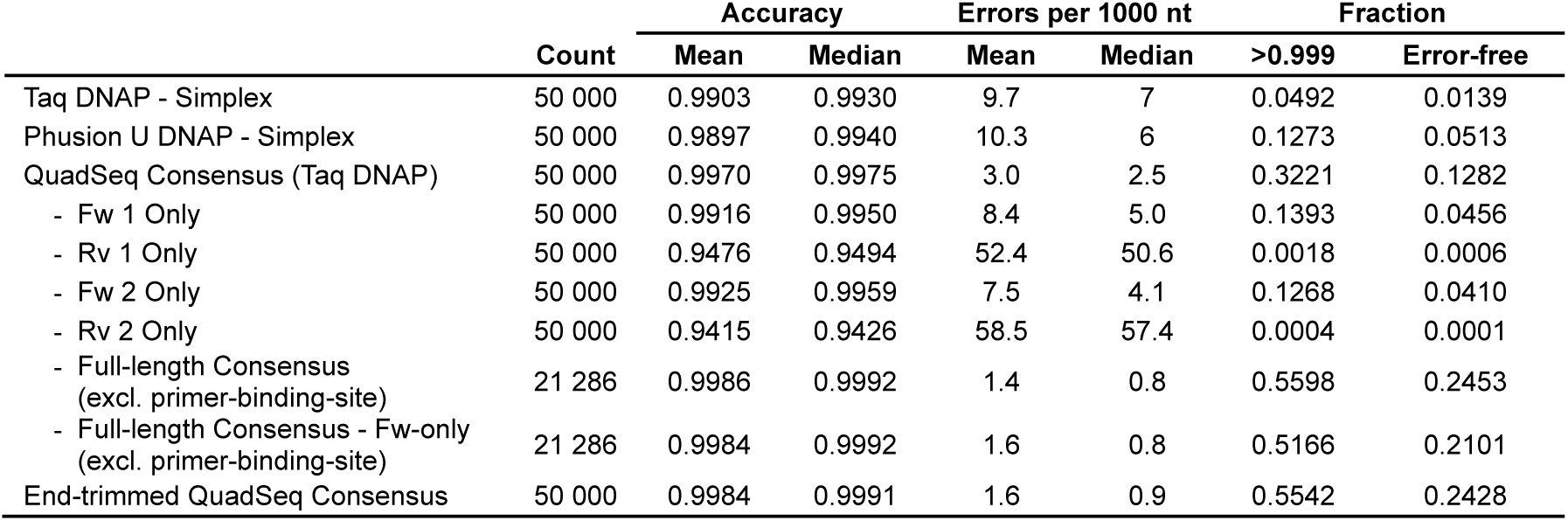
Nanopore sequencing accuracy of QuadSeq duplicated duplexes compared to simplex reads.

|  |  | Accuracy |  | Errors per 1000 nt |  | Fraction |  |
|---|---|---|---|---|---|---|---|
|  | Count | Mean | Median | Mean | Median | >0.999 | Error-free |
| Taq DNAP - Simplex | 50 000 | 0.9903 | 0.9930 | 9.7 | 7 | 0.0492 | 0.0139 |
| Phusion U DNAP - Simplex | 50 000 | 0.9897 | 0.9940 | 10.3 | 6 | 0.1273 | 0.0513 |
| QuadSeq Consensus (Taq DNAP) | 50 000 | 0.9970 | 0.9975 | 3.0 | 2.5 | 0.3221 | 0.1282 |
| - Fw 1 Only | 50 000 | 0.9916 | 0.9950 | 8.4 | 5.0 | 0.1393 | 0.0456 |
| - Rv 1 Only | 50 000 | 0.9476 | 0.9494 | 52.4 | 50.6 | 0.0018 | 0.0006 |
| - Fw 2 Only | 50 000 | 0.9925 | 0.9959 | 7.5 | 4.1 | 0.1268 | 0.0410 |
| - Rv 2 Only | 50 000 | 0.9415 | 0.9426 | 58.5 | 57.4 | 0.0004 | 0.0001 |
| - Full-length Consensus<br>(excl. primer-binding-site) | 21 286 | 0.9986 | 0.9992 | 1.4 | 0.8 | 0.5598 | 0.2453 |
| - Full-length Consensus - Fw-only<br>(excl. primer-binding-site) | 21 286 | 0.9984 | 0.9992 | 1.6 | 0.8 | 0.5166 | 0.2101 |
| End-trimmed QuadSeq Consensus | 50 000 | 0.9984 | 0.9991 | 1.6 | 0.9 | 0.5542 | 0.2428 |

### Full-length duplicated-duplex consensus sequences have a median accuracy of >Q30

The DNA-gap that is created when the USER III enzyme mixture cleaves the deoxyuridine base located in the forward QuadSeq primer enables a DNAP-mediated strand-displacement and DNA sequence duplication. However, because this cleavage occurs within the primer’s binding site, most of the forward primer sequence is absent from both the first and the last QuadSeq DNA copies, i.e. the first forward DNA strand and the second reverse strand (Fig. 2A). Previous studies have shown that nanopore reads have a lower basecalling accuracy in both the beginning and end of each sequencing read^24^, which we also observe, as the first forward DNA strand within the QuadSeq duplicated duplex has a lower median accuracy compared to the second forward DNA strand (Table 1). Thus, as the terminal ends of each nanopore read has a low accuracy, and the terminal ends of each QuadSeq consensus sequence is only derived from one high-fidelity forward strand and one low-fidelity reverse strand, i.e. the second forward DNA strand and the first reverse strand, we speculated that all QuadSeq consensus sequences would also have a much lower fidelity at the beginning of their sequences. To investigate this, we extracted only full-length QuadSeq duplicated duplex sequences and trimmed away both QuadSeq primer annealing sequences. We then generated a consensus sequence from these trimmed and filtered full-length fragments, ensuring that all 1177 nt were covered by all four fragments (Forward-Reverse-Forward-Reverse). This ensured that each consensus-derived nucleotide was selected from two high-fidelity and two low-fidelity sequences. The median accuracy of these trimmed and filtered full-length QuadSeq duplicated duplexes was 0.9992 (Q31), corresponding to only 0.8 errors per 1000 nt (Fig. 3E, Table 1). The fraction of reads with an accuracy greater than 0.999 (Q30) was 0.5598, and the fraction of error-free amplicons was 0.2453 (Table 1). Thus, demonstrating that QuadSeq consensus generation requires at least two high-fidelity DNA sequences and that full-length QuadSeq duplicated-duplexes can enable a nanopore sequencing fidelity that is approaching the PacBio Revio long-read sequencing platform (0.9995/Q33)^28^, while simultaneously retaining many of the advantages intrinsic to ONT sequencing, such as portability and cost.

### The reverse low-fidelity DNA copies in each duplicated-duplex can provide conditional error-corrections

Because we observed that both high-fidelity DNA copies encoded in the QuadSeq duplicated-duplex was required for high-fidelity nanopore sequencing, we next sought to quantify the beneficial effect that the two low-fidelity reverse DNA strands have on the QuadSeq consensus sequence. We repeated our previous consensus generation using full-length QuadSeq duplicated duplexes with the primer binding sites excluded, but now only used the two high-fidelity forward DNA strands to generate a consensus sequence. From this analysis, we observed that the median accuracy of full-length QuadSeq duplicated duplexes containing only the two forward DNA strands was also 0.9992 (Q31), corresponding to 0.8 errors per 1000 nt (Fig. 3E, Table 1). But the fraction of reads with an accuracy greater than 0.999 (Q30) was only 0.5166, and the fraction of error-free amplicons were only 0.2101 (Table 1). This quantification showed that the two low-fidelity reverse DNA strands in the QuadSeq duplicated-duplex consensus generation can increase the fraction of high-fidelity reads (>Q30) by almost 8%, and the fraction of error-free amplicons by almost 17% (Table 1). Thus, we observed that high-fidelity sequencing information was primarily derived from the two forward QuadSeq DNA strands but that the two low-fidelity reverse strands could provide a limited and modest error correction. These rare error correction events should only occur when the two forward DNA strands contain contradictory information and when both Phred quality scores are very similar. In these instances, the two reverse strands could act as tiebreakers, allowing for a minor, but beneficial, increase in nanopore sequencing accuracy.

### An end-trimming Nextflow pipeline for generating high-fidelity intramolecular consensus sequences from QuadSeq nanopore reads

By using all our previous learnings for creating intramolecular consensus sequences from QuadSeq duplicated-duplex DNA molecules, we next designed an improved Nextflow pipeline (DSL2) that generates high-fidelity single-read consensus sequences from ONT-basecalled QuadSeq FASTQ reads. The workflow of this Nextflow pipeline is as follows: **1)** ONT FASTQ reads are filtered based on user-defined read lengths (±2%). The expected read length of a fully formed QuadSeq duplicated-duplex molecule is 4 times the expected size of the amplicon + 168 nt (3 TelN-sites, each 56 nt in length). **2)** Fully formed duplicated duplex-reads are then split into four individual fragments/amplicons at each of their amplicon-connecting TelN-sites. **3)** All four extracted fragments/amplicons are trimmed by 30 nt at both the 5’ and 3’ ends to remove corrupting terminal ends. **4)** By then using gap penalties optimized for QuadSeq fragments (Table S4), a high-fidelity consensus sequence is generated from the DNA region that is covered by all four fragments by multiple sequence alignment (MSA). This consensus generation is based on a log-likelihood approach to select the base, or gap, with the highest quality-weighted log-likelihood based on the ONT Phred quality scores. This pipeline also implements a streaming architecture that processes ONT sequencing reads in batches, enabling real-time analysis and visualization during a nanopore sequencing run.

Next, we used our new pipeline to process QuadSeq ONT FASTQ reads to generate high-fidelity intramolecular consensus sequences. Because our QuadSeq pipeline uses a static-length end-trimming and an MSA consensus generation that is strictly based on all four amplicon copies (2 forward and 2 reverse), these high-fidelity consensus sequences are not identical in length. Yet this approach does ensure that the QuadSeq consensus sequences that are obtained have the highest possible accuracy. Similar to our previous analyses using filtered full-length QuadSeq fragments (Table 1), we could also observe that statically trimmed QuadSeq consensus sequences reached a very similar median accuracy (0.9991/Q30.5), corresponding to only 0.9 errors per 1000 nt (Fig. 3F, Table 1). The fraction of reads with an accuracy greater than 0.999 (Q30) was 0.5542, and the fraction of error-free amplicons was 0.2428 (Table 1). This data clearly demonstrate that QuadSeq-synthesized duplicated DNA duplexes can be utilized to generate high-fidelity single-read consensus sequences, which in the future could enable a ONT use even for difficult-to-sequence amplicons and complex and high-diversity DNA libraires.

### Full-length 16S rRNA QuadSeq metagenomic sequencing

One well-known ONT use case that can be difficult to undertake and interpret due to the high error-rate of ONT simplex sequencing is 16S rRNA metabarcoding^2, 3^. During this sequencing, the 16S rRNA gene, either the full-length gene, or different 16S rRNA variable regions, are amplified from a complex microbial community to identify and classify bacteria from an environmental or clinical sample without first growing them in the lab. Because bacterial 16S rRNA genes often have very similar sequences, high ONT error rates can be determinantal for accurately identifying closely related bacterial species^2, 3^. Thus, we sought to determine whether QuadSeq ONT sequencing could be utilized for accurate bacterial species identification of full-length 16S rRNA amplicons. To achieve this, we amplified the 16S rRNA gene from a consortium of 8 pre-purified bacterial genomes from ZymoBIOMICS, including; *Listeria monocytogenes, Pseudomonas aeruginosa, Bacillus subtilis, Escherichia coli, Salmonella enterica, Lactobacillus fermentum, Enterococcus faecalis, Staphylococcus aureus*, and 2 pre-purified fungal genomes, including; *Saccharomyces cerevisiae* and *Cryptococcus neoformans*, using the standardized 27F and 1492R amplifications primers and Taq DNAP^46^. To enable a 16S QuadSeq duplicated-duplex-synthesis, we also amplified the 16S rRNA loci using QuadSeq-enabled 27F and 1492R primers, which encoded the 56 nt TelN-site and 5 additional non-base-pairing nucleotides (Fig. 2F). After a QuadSeq DNA-synthesis, the 16S duplicated duplexes were ONT sequenced and each QuadSeq read was used to generate a high-fidelity intramolecular consensus sequence (Fig. 2F, Table 2). From this experiment, we could observe that the median accuracy of end-trimmed (30 nt) simplex reads of Taq-DNAP-synthesized full-length 16S rRNA genes were 0.9952, corresponding to 7.2 errors per 16S amplicon (estimated at 1500 nt), and the fraction of 16S simplex reads with an accuracy greater than 0.999 (Q30) was 0.1055 (Fig. 4A, Table 2). In contrast, end-trimmed QuadSeq consensus sequences of full-length 16S rRNA amplicons hade a median accuracy of 0.9986, corresponding to 2.1 errors per 16S rRNA amplicon, and the fraction of high-fidelity reads (>Q30) were 0.3474 (Fig. 4A, Table 2), demonstrating that QuadSeq consensus sequences can decrease the median error-rate of 16S rRNA amplicons by more than 3-fold, and the fraction of high-fidelity amplicons (>Q30) by almost 4-fold (Table 2), thus increasing both the accuracy and reliability of 16S rRNA metabarcoding.

**Figure 4.**
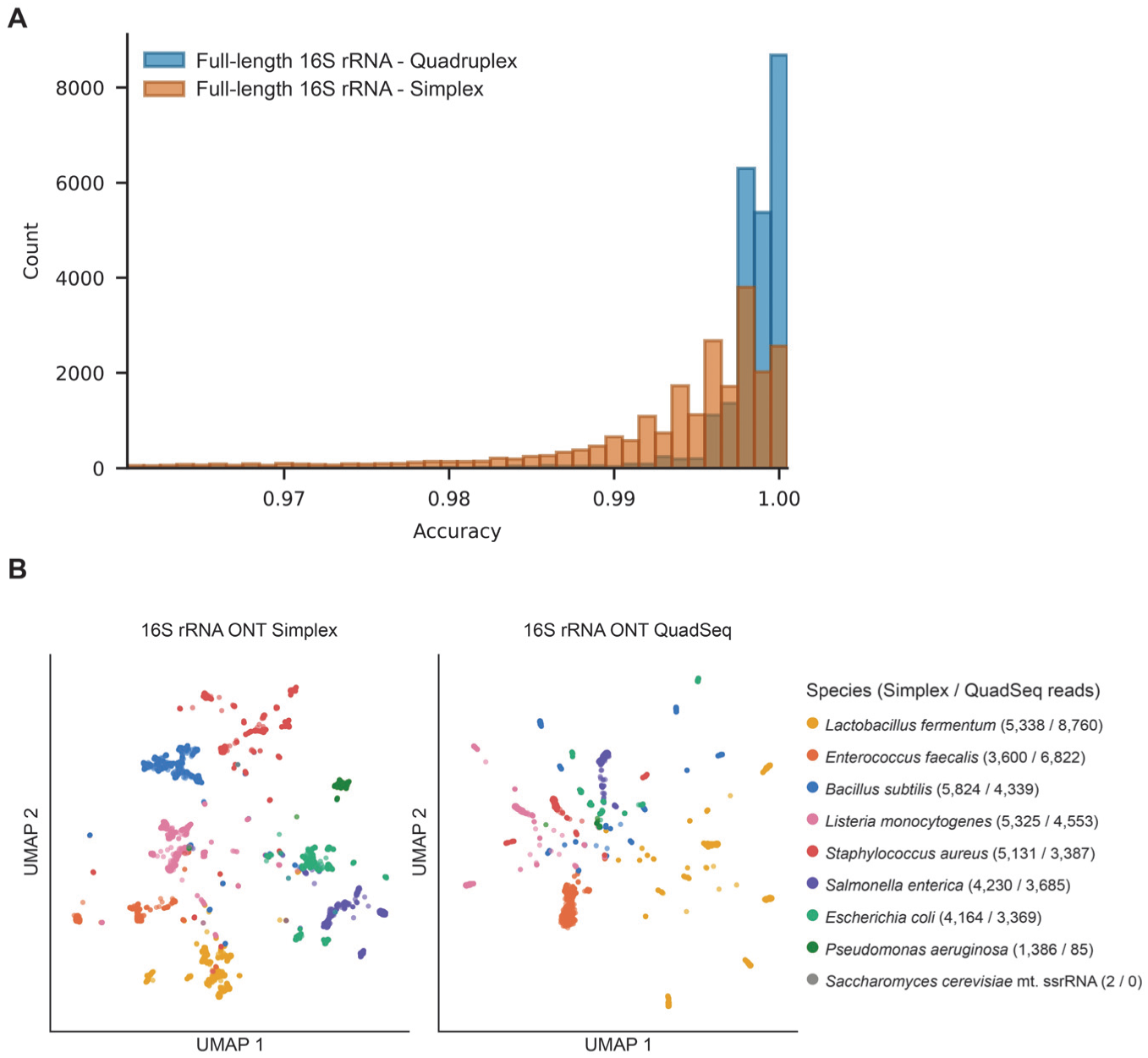
QuadSeq ONT sequencing facilities better species level resolution during 16S rRNA metabarcoding. **A)** QuadSeq intramolecular consensus sequences from end-trimmed full-length 16S rRNA amplicons can reach a median single-amplicon accuracy of 0.9986 (Q28.5). **B)** A two-dimension UMAP of the relative frequencies of both simplex and QuadSeq 16S rRNA consensus sequences represented by their canonical 6-mer frequencies, with each k-mer merged with its reverse complement (2080 features). Profiles were reduced to 50 principal components and embedded in two dimensions with UMAP. The two datasets were embedded separately, so their axes are not comparable. Datapoints are colored by the species that best match the 26 unique small subunit rRNA loci found in the ZymoBIOMICS microbial community DNA library.

**Table 2.**
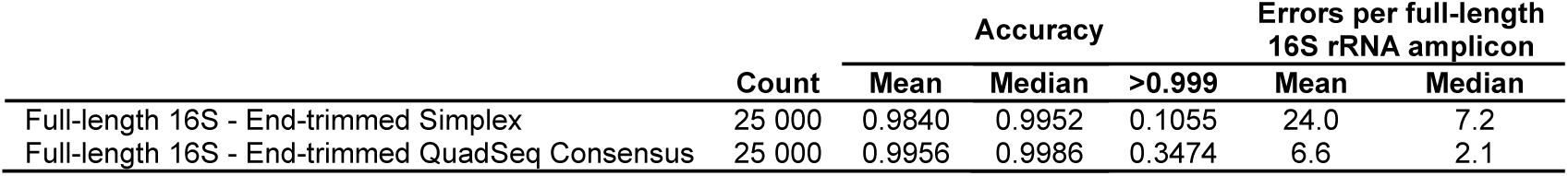
Full-length 16S rRNA ONT QuadSeq amplicon sequencing.

### QuadSeq metagenomic sequencing minimizes overall false-positive species detection

We observed that QuadSeq 16S rRNA metagenomic sequencing decreased ONT sequencing errors from 7.2 to 2.1 errors per full-length 16S amplicon. Thus, we next sought to investigate if this increased accuracy would also translate to an improved genus- and species-level resolution. Both high-fidelity 16S rRNA QuadSeq consensus sequences and 16S rRNA simplex reads were mapped against the ZymoBIOMICS small subunit rRNA reference sequence library (containing 26 unique rRNA reference sequences from 8 bacterial genomes, and 2 fungal genomes) that were used as template for the 16S metagenomic PCR (Fig. 2F, 4B). Their relative relationships were then visualized by a uniform manifold approximation and projection for dimension reduction (UMAP) analysis, demonstrating that a single-read error-correction facilitated by full-length 16S rRNA QuadSeq ONT sequencing can ensure true positive detection of low-frequent species, while minimizing an overall false-positive species detection (Fig. 4B). Yet we could also observe that the addition of the TelN-site (56 nt) to the 27F and 1492R primer sequences changed the relative DNA-amplification rate of the *P. aeruginosa* 16S rRNA loci (Fig. 4B). QuadSeq 16S amplicons mapping to the 16S loci of *P. aeruginosa* was almost nonexistent among the generated QuadSeq consensus sequences. It is likely that the addition of the TelN-site(s) interferes with the annealing of the 27F and/or 1492R primers to the 16S rRNA loci of *P. aeruginosa*, which could cause a selective attenuation of this specific 16S amplicon from the QuadSeq metagenomic PCR library. To resolve this issue in the future, we suggest that alternative 16S metabarcoding primers are empirically evaluated on a case-by-case basis for their QuadSeq compatibility before any extensive metagenomic sequencing project is undertaken.

### A model for generating estimated Phred quality scores for QuadSeq consensus sequences

A common strategy during most post-sequencing analysis is to filter reads by their average Phred quality scores and then only use high-fidelity reads, often >Q30, for any downstream analysis. This ensures that any conclusion(s) made from the data is both reliable and robust. QuadSeq high-fidelity duplicated-duplex consensus sequences have higher accuracy than nanopore simplex reads, but not all QuadSeq consensus sequences are suited for all downstream analyses. Thus, to enable a Phred quality score parsing of QuadSeq duplicated-duplex consensus sequences, we also developed a model that derived estimated consensus quality scores directly from the Phred scores encoded in each basecalled nanopore FASTQ read generated by the ONT algorithm (Fig. 3F). Using our previous learnings, we designed this model to use the arithmetic mean per base of the two fragments with the highest Phred scores in each duplicated duplex-read. For the majority of nanopore sequencing reads, this meant taking the mean Phred score from the two forward high-fidelity DNA strands (Fig. 3F). We then mapped the average estimated Phred score vs. the observed Phred score for each generated consensus sequence (Fig. 3F). This model accurately generated estimated Phred scores that closely correlated with the observed Phred scores. By combining this model and our new custom QuadSeq Nextflow pipeline, it is possible to generate Phred-score-encoded QuadSeq duplicated-duplex consensus sequences that can be filtered using conventional post-sequencing analysis cut-offs, such as only parsing reads >Q30. This model should significantly improve the utility of the QuadSeq technology and enable an easy high-fidelity nanopore amplicon sequencing and bioinformatic analysis that can effectively compete with the PacBio Revio platform on most relevant metrics.

## Discussion

Long-read ONT sequencing is a highly versatile and portable technology that is constrained by relatively high error rates compared to competing platforms such as PacBio Revio or Vega^1^. Here we describe QuadSeq, a three-step DNA-synthesis technology that can convert linear amplicons into duplicated DNA duplexes encoding two forward and two reverse strand copies, thereby enabling high-fidelity real-time nanopore amplicon sequencing. Unlike other strategies aimed at increasing the accuracy of nanopore sequencing, such as UMI-barcoding^1, 15^, QuadSeq derives an intramolecular consensus from each individual ONT read independently, enabling high-fidelity sequencing in real time when coupled with our custom Nextflow pipeline, which can processes, align and generate consensus sequences in batches.

QuadSeq duplicated duplexes are end-point molecules, i.e. the DNA-synthesis/DNA-duplication proceeds only until each molecule is fully formed. Thus, because every successfully synthesized duplicated duplex has a predictable and uniform molecular structure, a size-dependent DNA-purification and targeted sequencing is possible. This contrasts with conventional ONT duplex sequencing, which is often limited to generating high-fidelity consensus sequences from less than 10% of all available ONT reads^21, 22^. The ability to easily titrate the USER III enzymes during the QuadSeq reaction also enables a staggered DNA-synthesis, which can minimize the formation of DNA chimeras, as only a few DNA molecules are actively synthesized at any given time. In addition, the QuadSeq reaction time can be extended for samples with difficult-to-amplify sequences to increase the chance that all DNA molecules are fully formed during the reaction. This can minimize DNA-amplification biases and is a feature of the QuadSeq technology that is in stark contrast to both the RCA- and MDA-dependent amplification strategies, which are constrained by high levels of DNA-inversions, rearrangements and chimera formations, as well as amplification biases^30–32^.

Nanopore sequencing of inverted-repeat DNA-molecules, such as physically-linked DNA duplexes, is known to impair the basecalling fidelity of the reverse/complementary DNA strand because of a DNA-reannealing *in trans*^24, 25^. However, the unique molecular structure of the QuadSeq duplicated duplex partially overcomes this inhibitory DNA-structure. During nanopore sequencing, each duplicated DNA duplex transitions through a nanopore in four distinct phases. During the first and third phase, i.e. the transition of the two forward DNA strands, each fragment can be basecalled with a high accuracy, whereas during the second and fourth phase, i.e. the transition of the two reverse DNA strands, the basecalling-fidelity is corrupted by an inhibitory DNA secondary structure formation. Consequently, each QuadSeq duplicated-duplex is comprised of two high-fidelity and two low-fidelity DNA copies of the original amplicon. Yet, when all four copies are used to generate an intramolecular consensus sequence, the median single-amplicon accuracy of each duplicated-duplex can reach 0.9992 (Q31). The majority of this error-correction power is derived from the two high-fidelity forward DNA strands, but the two low-fidelity reverse DNA strands can provide conditional error-corrections by acting as consensus-tiebreakers. Thus, nanopore sequencing of QuadSeq duplicated duplexes can partially overcome most constraints that have previously corrupted ONT sequencing of physically linked DNA duplexes (Fig. 1C). Furthermore, the nanopore accuracy that is achievable when deriving a consensus sequence from two high-fidelity and two low-fidelity DNA copies can also rival the reported accuracy of the PacBio Revio platform (0.9995/>Q33)^28^, while simultaneously retaining many of the advantages intrinsic to ONT sequencing, such as portability and cost.

One of the most consequential findings of this study is that the primary bottleneck for accurate nanopore amplicon sequencing is not the mutation rate of the initial PCR-amplification, but the fidelity of the nanopore sequencing. End-trimmed QuadSeq consensus sequences generated from amplicons synthesized by the low-fidelity Taq DNAP had a higher median nanopore accuracy (0.9991/Q30.5) than simplex reads of amplicons synthesized by the high-fidelity Phusion U DNAP (0.9940/≈Q22). In the future, this result could enable a direct PCR-amplification of high-value amplicons-of-interests found in both crude and dirty samples, such as whole blood or soil extracts, without sacrificing sequencing fidelity. I.e. the initial PCR-amplification can be performed using a robust, but low-fidelity DNAP, without requiring an additional DNA-purification step that could introduce a DNA-enrichment bias^34–37^.

By also using our model for estimating consensus Phred scores directly from the duplicated-duplex nanopore FASTQ sequences, each QuadSeq consensus sequence can be filtered using conventional post-sequencing analysis cut-offs, such as only parsing reads with an Phred score above Q30. This can restrict the downstream bioinformatic analysis to high-fidelity reads, which can further increase the accurate genotyping of rare DNA variants. Together with our streaming Nextflow consensus-generating pipeline, this should facilitate a straightforward integration of QuadSeq sequencing data into most pre-existing bioinformatic workflows.

Yet, one limitation of the QuadSeq technology is that each duplicated-duplex amplicon occupies four times the read length of a simplex read, while primarily only the two high-fidelity forward DNA copies, i.e. half the length of the total read, contributes to the consensus error-correction. In theory, this means that the QuadSeq duplicated duplexes decrease the effective throughput of each ONT sequencing run by approximately half, compared to when using either an RCA, MDA or UMI-barcoding approaches. Though, this is difficult to precisely quantify, because the relationship between read length and flow cell output is not linear^47, 48^. For example, Baslan *et al.* observed that when they sequenced purified DNA from a SK-BR-3 breast cancer cell line that was fragmented to an average length of either 400 or 10 000 bp, one nanopore flow cell (MinION R9.4.1) could read 6 million molecules from the small-fragment (400 nt) library, but only 1 million DNA molecules from the large-fragment (10 000 nt) library^47^. Yet, because each of these DNA molecules were approximately 25-times bigger, the sequencing data output per flow cell was still more than 4-fold greater for the large-fragment library than for the small-fragment library. Thus, for short amplicons (1500 nt) that would be sequenced as an approximately 6000 nt long duplicated-duplex, the increased read-length of each QuadSeq duplicated DNA duplex would not necessarily reduce the total sequencing data output, as each nanopore flow cell could potentially process a similar number of medium-length (6000 nt) amplicons compared to short-length (1500 nt) amplicons. Consequently, QuadSeq sequencing of small amplicons would simply add additional sequencing data, which could be used for high-fidelity consensus generation, without necessarily limiting the total data output of each flow cell. However, this hypothesis needs to be tested empirically for each individual amplicon-of-interest and for many different amplicon lengths.

## Data availability

The sequencing data for this study have been deposited in the National Center for Biotechnology Information (NCBI) under bio project accession: PRJNA1556708. Source data are provided with this paper.

## Code availability

The QuadSeq pipeline (v0.1.0) and the QuadSeq nanopore sequencing data analysis are available from the Novo Nordisk Foundation Biotechnology Research Institute for the Green Transition (BRIGHT) GitHub repository for this project (https://github.com/BRIGHT-CDA/quadseq).

## Author contributions

**Marcus Wäneskog**: Conceptualization; Designed research; Performed research; Analyzed data; Funding acquisition; Wrote the original draft, review(s) and editing. **Knud Nor Nielsen**: Performed research; Analyzed data; Wrote review(s) and editing. **Achille Julienne**: Analyzed data. **Carlos G. Acevedo-Rocha**: Project administration; Resources; Wrote review(s) and editing. **Emre Özdemir**: Project administration; Resources. **Emil Damgaard Jensen**: Project administration; Resources; Funding acquisition; Wrote review(s) and editing.

## Material and methods

### Vector constructs

The plasmids used in this study are listed in Table S1.

### Oligonucleotides

The oligos and PCR primers used in this study are listed in Table S2. Oligos were purchased from Integrated DNA Technologies (IDT) either as desalted or polyacrylamide gel electrophoresis (PAGE) purified oligonucleotides and shipped lyophilized. QuadSeq-enabled forward primers were modified to contain a deoxyuridine base close to the 3’-end, but not directly at the 3’-end of the oligo, by replacing a naturally occurring thymine base in the primer annealing sequence. To facilitate easy downstream ONT-adapter ligations, PCR primers used to amplify control (Simplex) amplicons were synthesized with a phosphate group added to the 5’-end.

### QuadSeq PCR-amplifications

PCR reactions were performed using QuadSeq-enabled forward and reverse primers, which encoding TelN-sites in their 5’-ends (Table S2), using either the Phusion U hot start PCR master mix (Thermo Scientific), DreamTaq hot start PCR master mix (Thermo Scientific), Red Taq PCR master mix (VWR Chemicals), or OneTaq hot start PCR master mix (New England Biolabs). PCR cycling protocols were carried out according to the manufacturers’ recommendations using 10 ng of a plasmid DNA template (Table S1) for each 50 µl reaction, and 1 µM of each QuadSeq primer (final concentration) (Table S2).

### QuadSeq 16S rRNA gene PCR amplifications

PCR reactions were performed using QuadSeq-enabled 27F and 1492R forward and reverse primers^46^ (Table S2), using the Red Taq PCR master mix (VWR Chemicals). PCR cycling protocols were carried out according to the manufacturers’ recommendations using 100 ng of ZymoBIOMICS microbial community DNA standard (Zymo Research) as the template for each 50 µl reaction, and 1 µM of each QuadSeq primer (final concentration) (Table S2). The ZymoBIOMICS DNA standard contains 8 purified bacterial and 2 purified fungal genomes, including *Listeria monocytogenes, Pseudomonas aeruginosa, Bacillus subtilis, Escherichia coli, Salmonella enterica, Lactobacillus fermentum, Enterococcus faecalis, Staphylococcus aureus, Saccharomyces cerevisiae*, and *Cryptococcus neoformans*.

### TelN-mediated DNA-end sealing

Linear DNA amplicons encoding TelN-sites at each terminal end was suspended in a 1x ThermoPol reaction buffer (New England Biolabs), containing 20 mM Tris-HCl, 10 mM (NH_4_)_2_SO_4_, 10 mM KCl, 2 mM MgSO_4_, 0.1% Triton X-100 at pH 8.8, at a final concentration of 20 µg/ml (approx. 25 pmol/ml). The TelN Protelomerase (New England Biolabs) was then added at a final concentration of 250 units/ml and the mixture was incubated for 2 hours at 30 °C. If the TelN-site-encoded DNA amplicons were PCR-amplified using non-PAGE-purified QuadSeq primers, then 100 units/ml of Exonuclease III (New England Biolabs) and 60 units/ml of Thermolabile Exonuclease I (New England Biolabs), or 20 units/ml of Exonuclease VIII (New England Biolabs) and 60 units/ml of Thermolabile Exonuclease I (New England Biolabs), was next added to the ThermoPol reaction mixture (final concentration), followed by an additional 1 hour incubation at 37 °C. The resulting covalently closed DNA amplicons were then purified by DNA-binding silica column.

### QuadSeq duplicated-duplex DNA synthesis

A detailed protocol description of the QuadSeq workflow (Table S3) can be found in the supplemental online material for this article.

### Agarose gel electrophoresis

25 ng of each amplicon and QuadSeq duplicated-duplex DNA sample was separated and visualized in-gel by agarose gel electrophoresis, made by mixing and solidifying 1% melted agarose in 1x TAE-buffer (40 mM tris(hydroxymethyl)aminomethane, 20 mM acetic acid and 1 mM ethylene diamine tetraacetic acid) containing SYBR Gold nucleic acid stain (Thermo Scientific), diluted 10 000-fold, according to manufacturer’s instructions.

### ONT sequencing

Either 800 ng of DNA-binding-column-purified QuadSeq duplicated-duplex DNA (approx. 400 fmol), or 400 ng of DNA-binding-column-purified linear DNA controls (approx. 400 fmol) was used as the template for each ONT library preparation. Sequencing libraries were prepared using the SQK-LSK114 (kit V14) ligation sequencing kit, following the manufacturer’s protocol, except that a higher then recommended DNA input (400 fmol) was used per reaction. This was to account for one covalently closed and inaccessible DNA-end in the QuadSeq duplicated-duplex molecule. DNA samples were washed with ONT long fragment buffer to enrich for longer (i.e. full-length) duplicated duplex DNA fragments. Samples were sequenced on a MinION Mk1B device using R10.4.1 (FLO-MIN114) flow cells, following the manufacturer’s instructions. Reads were basecalled with Dorado v0.9.6 in super-accuracy mode with model dna_r10.4.1_e8.2_400bps_sup@v5.0.0.

Each DNA library was sequenced as a stand-alone sample, i.e. no sample multiplexing was performed, except for the Taq vs. Phusion U Simplex control. For these DNA samples, Taq-amplified DNA fragments were pooled with Phusion-U-amplified DNA fragments and sequenced as one ONT library. To distinguish these two samples, Taq-amplified DNA fragments were tagged with terminal TelN-sites, while Phusion U DNA fragments were amplified without TelN-sites. This enabled a post-sequencing parsing of both DNA populations/samples.

### QuadSeq Nextflow pipeline design

The QuadSeq pipeline (v0.1.0) is implemented in Nextflow^49^ (v26.04 or later) and converts each nanopore read of a duplicated DNA duplex into one intramolecular consensus sequence. All steps are run in Docker or Singularity containers, or a conda environment, using Python 3.10 and Biopython^50^. FASTQ files are split into batches of reads (1000 by default) that are processed independently, so that results are accumulated and reported progressively. In real-time mode, the pipeline monitors the sequencer output folder, processes each FASTQ file as it is written, and updates an HTML report after every batch.

#### Length filtering and fragmentation

A complete QuadSeq molecule has a predictable length: four amplicon copies joined by three 56 nt linker motifs. Reads were therefore first filtered on length. By default, the pipeline keeps reads within ±2% of the length peak, the read length L whose window 0.98L-1.02L contains the most reads, recomputed as reads accumulate. A fixed window can also be set, or the filter can be disabled.

Linker motifs (5′-TATCAGCACACAATNNNCCATTATACGCGCGTATAATGGNNNATTGTGTGCTGATA-3′, N set to A) were located on both strands with blastn (BLAST+ v2.17.0^51^; word size 5, reward 1, penalty −2, gap open 5, gap extension 2, no DUST filtering), keeping hits with a raw score above 16. As the motif is palindromic, overlapping hits on opposite strands were reduced to the forward hit. Reads were cut at each hit, and the motif sequences were removed. Fragments outside a user-defined amplicon length range were discarded, and 30 nt (default) were trimmed from both ends of each remaining fragment. Molecules were classified by their number of fragments as quadruplex (4), triplex (3), duplex (2) or simplex (1). Reads with more than three motifs (concatemers) were excluded. By default, only quadruplex molecules are aligned, though triplex, or all molecule types can be included.

#### Alignment and consensus calling

The fragments of each molecule were aligned with MAFFT^52^ (v7.526) E-INS-i (--genafpair --maxiterate 1000 --adjustdirection), using gap parameters optimized for QuadSeq (--op 0.637 --ep 0 --lop −0.775 --lep 0.235 --lexp 0) (Table S4). The alignment was restricted to the columns spanned by all fragments. Each gap was assigned the mean Phred quality^53^ of the flanking bases in its fragment. In each column, the consensus character b ε (A, C, G, T, -) was the one maximizing

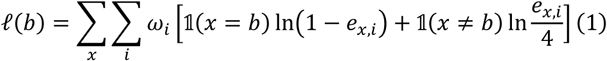

where eₓ,ᵢ = 10^(−Q/10) is the error probability of the i-th best observation (i = 0, 1, …) of character x, ranked by quality. The weights ωo = 1 and ωᵢ = w/(i + 1) for i ≥ 1, with w = 2, reduce the influence of repeated observations of the same character. Columns called as gaps were omitted. Each consensus base was given the quality of the mean error probability of its two best supporting observations. Simplex molecules were reported as their single trimmed fragment.

#### Optimization of alignment parameters

The five MAFFT gap parameters and w were optimized jointly by Markov chain Monte Carlo sampling with simulated annealing^54, 55^ (Metropolis acceptance; initial temperature 1, cooling factor 0.95 per step). The objective was the mean observed Q-score of the consensus sequences of 883 molecules of known sequence (611 quadruplex, 272 triplex), computed after mapping to the reference with minimap2 v2.28^56, 57^. Across 31 chains, 662 parameter sets were evaluated, and the best set (mean Q-score 25.8) became the pipeline default (Table S4).

#### Consensus accuracy

Optionally, consensus sequences are added with –add^58^ to a cached MAFFT E-INS-i alignment of user-supplied reference sequences. For each consensus–reference pair, terminal columns containing a gap and then terminal mismatches were trimmed. Accuracy was calculated as M/(M + X + I + D), where M, X, I and D are matches, mismatches, insertions and deletions, respectively. The observed Q-score is −10 log4o(1 − accuracy), set to 60 for perfect matches. Each consensus was assigned to the reference with the highest accuracy, with ties broken by the number of matches.

#### Taxonomic classification

Optionally, consensus sequences are classified to species level with EPI2ME wf-16s (v1.6.1; https://github.com/epi2me-labs/wf-16s) using default parameters. wf-16s aligns reads with minimap2 v2.28 to the NCBI 16S/18S targeted loci database^59^ and assigns NCBI Taxonomy^60^ lineages. Species counts were accumulated across batches and shown in the live report.

### Bioinformatic analyses

Four ONT datasets were analyzed. **catR simplex**: catR amplified with Taq DNAP, whose amplicons carried TelN sites, or with Phusion U DNAP, whose amplicons did not, sequenced together as standard simplex reads. **catR QuadSeq**: QuadSeq duplicated DNA duplexes made from Taq-amplified catR. **16S QuadSeq**: QuadSeq duplicated DNA duplexes of 16S rRNA gene amplicons from the ZymoBIOMICS Microbial Community DNA Standard (Zymo Research). **16S simplex**: the same 16S amplicons sequenced as standard simplex reads. All DNA libraires were sequenced on a MinION Mk1B device using R10.4.1 (FLO-MIN114) flow cells, and basecalled with Dorado super-accuracy model, sup@v5.0.0.

#### catR simplex

Reads were separated by the presence of the TelN motif. Phusion U reads were identified by their lack of a TelN-motif, 1,219-1,244 nt long, while Taq reads were identified by the presence of a TelN-motif at both ends, 1,215–1,233 nt long. We sampled 60 000 reads of each population at random, aligned each read to the catR template with MAFFT (--6merpair --adjustdirection) and calculated accuracies same as for QuadSeq consensus sequences. Random subsets of 50 438 Taq reads served as the simplex comparison for catR QuadSeq.

#### catR QuadSeq

Consensus accuracies were measured against the catR template. For Fig. 3B-C, molecules were kept if all four fragments were 1163-1284 nt (±5% around 1224 nt), giving 51 093 molecules for fragment accuracies and 50 438 for consensus accuracy. For Fig. 3E-F, we selected molecules with a read length of 4950-5100 nt, around the expected 5060 nt, corresponding to four amplicons and three 56 nt TelN sites. Fig. 3E uses 22 672 of these in which all four fragments contained the full-length amplicon sequence, minus the primer-binding sites, with consensus sequences called either from all four fragments or from the two forward fragments re-aligned on their own. Fig. 3F uses a random sample of 50 000 reads, processed with the QuadSeq pipeline described above (Nextflow v26.04.6; batches of 5000 reads; fragments 1101-1345 nt, ±10% around the 1223 nt template).

#### 16S QuadSeq and 16S simplex

Both datasets were processed with the QuadSeq pipeline (Nextflow v26.04.6; batches of 5000 reads), with fragments filtered to 1464-1618 nt, ±5% around the 1541 nt median length of the bacterial 16S ZymoBIOMICS Microbial Community DNA Standard (Zymo Research) reference sequences, and with no whole-molecule length filter. Quadruplex consensus sequences were analyzed for 16S QuadSeq and single-trimmed fragments of 16S simplex reads. Each sequence was assigned to its best-matching reference among the 26 unique small subunit rRNA sequences of the ZymoBIOMICS reference set (refseq v3), which served as ground truth, and was classified with EPI2ME wf-16s.

#### Metagenomic UMAP

35 000 processed sequences from each 16S dataset were sampled at random. Each sequence was represented by its canonical 6-mer frequencies, with each k-mer merged with its reverse complement (2080 features). Profiles were reduced to 50 principal components with scikit-learn and embedded in two dimensions with UMAP (umap-learn; 1000 neighbours, minimum distance 0, Euclidean metric; random seed 42). The two datasets were embedded separately, so their axes are not comparable, and points were colored by best-matching reference species.

### Code

The QuadSeq pipeline v0.1.0 (https://github.com/BRIGHT-CDA/quadseq), was implemented in Nextflow v26.04.6 and Python v3.10.21 with Biopython v1.88, pandas v2.3.3, NumPy v2.2.6 and Plotly v7.0.0. Within the pipeline, linker motifs were located with BLAST+ v2.17.0 and fragments were aligned with MAFFT v7.526. Taxonomy was assigned with EPI2ME wf-16s v1.6.1 (run with Nextflow v24.10.5), which uses minimap2 v2.28-r1209. Accuracy analyses used Python v3.10.21 with pandas v2.3.3, NumPy v2.2.6 and Matplotlib v3.10.9. Other downstream analyses and figures used Python v3.12.14 with pandas v3.0.5, NumPy v2.5.2, SciPy v1.18.1, statsmodels v0.15.0, Matplotlib v3.11.1, Biopython v1.88 and edlib v1.3.9. Reference sequences were deduplicated with SeqKit v2.13.0. For the phylogeny, sequences were aligned with MAFFT v7.526 L-INS-i, and a maximum-likelihood tree was inferred with IQ-TREE v3.1.3 (ModelFinder Plus; 1000 ultrafast bootstrap and 1000 SH-aLRT replicates). k-mer profiles were embedded with scikit-learn v1.9.1 and umap-learn v0.5.12, in Python v3.12 with pandas v3.0.6, NumPy v2.5.4 and Plotly v7.1.0.

## References

1. Karst, S.M. et al. High-accuracy long-read amplicon sequences using unique molecular identifiers with Nanopore or PacBio sequencing. Nat Methods 18, 165–169 (2021).

2. Matsuo, Y. et al. Full-length 16S rRNA gene amplicon analysis of human gut microbiota using MinION nanopore sequencing confers species-level resolution. BMC Microbiol 21, 35 (2021).

3. Aja-Macaya, P. et al. Nanopore full length 16S rRNA gene sequencing increases species resolution in bacterial biomarker discovery. Sci Rep 15, 26486 (2025).

4. Lao, H.Y. et al. The clinical utility of Nanopore 16S rRNA gene sequencing for direct bacterial identification in normally sterile body fluids. Front Microbiol 14, 1324494 (2023).

5. Moon, J. et al. Rapid diagnosis of bacterial meningitis by nanopore 16S amplicon sequencing: A pilot study. Int J Med Microbiol 309, 151338 (2019).

6. Han, H.S., Ro, D.H., Chung, J., Kim, N. & Moon, J. Nanopore 16S amplicon sequencing enables rapid detection of pathogen in knee periprosthetic joint infection. Int J Med Microbiol 312, 151570 (2022).

7. Quick, J. et al. Multiplex PCR method for MinION and Illumina sequencing of Zika and other virus genomes directly from clinical samples. Nat Protoc 12, 1261–1276 (2017).

8. Cabibbe, A.M. et al. Application of Targeted Next-Generation Sequencing Assay on a Portable Sequencing Platform for Culture-Free Detection of Drug-Resistant Tuberculosis from Clinical Samples. J Clin Microbiol 58 (2020).

9. Liu, P.Y. et al. Comprehensive pathogen identification and antimicrobial resistance prediction from positive blood cultures using nanopore sequencing technology. Genome Med 16, 141 (2024).

10. Bloemen, B. et al. Overcoming challenges in metagenomic AMR surveillance with nanopore sequencing: a case study on fluoroquinolone resistance. Front Microbiol 16, 1614301 (2025).

11. Taxt, A.M., Avershina, E., Frye, S.A., Naseer, U. & Ahmad, R. Rapid identification of pathogens, antibiotic resistance genes and plasmids in blood cultures by nanopore sequencing. Sci Rep 10, 7622 (2020).

12. Matern, B.M. et al. Long-Read Nanopore Sequencing Validated for Human Leukocyte Antigen Class I Typing in Routine Diagnostics. J Mol Diagn 22, 912–919 (2020).

13. Mosbruger, T.L. et al. Utilizing nanopore sequencing technology for the rapid and comprehensive characterization of eleven HLA loci; addressing the need for deceased donor expedited HLA typing. Hum Immunol 81, 413–422 (2020).

14. Tan, D.Z.J. et al. Developing a Nanopore Sequencing Workflow for Protein Engineering Applications. ACS Synth Biol 12, 2041–2050 (2023).

15. Zurek, P.J., Knyphausen, P., Neufeld, K., Pushpanath, A. & Hollfelder, F. UMI-linked consensus sequencing enables phylogenetic analysis of directed evolution. Nat Commun 11, 6023 (2020).

16. Currin, A. et al. Highly multiplexed, fast and accurate nanopore sequencing for verification of synthetic DNA constructs and sequence libraries. Synth Biol (Oxf) 4, ysz025 (2019).

17. Long, Y. et al. LevSeq: Rapid Generation of Sequence-Function Data for Directed Evolution and Machine Learning. ACS Synth Biol 14, 230–238 (2025).

18. Khrenova, M.G. et al. Nanopore Sequencing for De Novo Bacterial Genome Assembly and Search for Single-Nucleotide Polymorphism. Int J Mol Sci 23 (2022).

19. Yu, F. et al. Alignment-free unique molecular identifier clustering suppresses sequencing errors for accurate detection of low-frequency DNA variants. Brief Bioinform 26 (2025).

20. Helsmoortel, M. et al. Oxford Nanopore enhanced accuracy of long-read amplicons applied to microbial whole-genome sequencing. Microbiol Spectr 14, e0285625 (2026).

21. Sanderson, N.D. et al. Comparison of R9.4.1/Kit10 and R10/Kit12 Oxford Nanopore flowcells and chemistries in bacterial genome reconstruction. Microb Genom 9 (2023).

22. Lerminiaux, N., Fakharuddin, K., Mulvey, M.R. & Mataseje, L. Do we still need Illumina sequencing data? Evaluating Oxford Nanopore Technologies R10.4.1 flow cells and the Rapid v14 library prep kit for Gram negative bacteria whole genome assemblies. Can J Microbiol 70, 178–189 (2024).

23. Jain, M., Olsen, H.E., Paten, B. & Akeson, M. The Oxford Nanopore MinION: delivery of nanopore sequencing to the genomics community. Genome Biol 17, 239 (2016).

24. Krishnakumar, R. et al. Systematic and stochastic influences on the performance of the MinION nanopore sequencer across a range of nucleotide bias. Sci Rep 8, 3159 (2018).

25. Spealman, P., Burrell, J. & Gresham, D. Inverted duplicate DNA sequences increase translocation rates through sequencing nanopores resulting in reduced base calling accuracy. Nucleic Acids Res 48, 4940–4945 (2020).

26. Li, C. et al. INC-Seq: accurate single molecule reads using nanopore sequencing. Gigascience 5, 34 (2016).

27. Volden, R. et al. Improving nanopore read accuracy with the R2C2 method enables the sequencing of highly multiplexed full-length single-cell cDNA. Proc Natl Acad Sci U S A 115, 9726–9731 (2018).

28. Wenger, A.M. et al. Accurate circular consensus long-read sequencing improves variant detection and assembly of a human genome. Nat Biotechnol 37, 1155–1162 (2019).

29. Oliynyk, R.T. & Church, G.M. Efficient modification and preparation of circular DNA for expression in cell culture. Commun Biol 5, 1393 (2022).

30. Lasken, R.S. & Stockwell, T.B. Mechanism of chimera formation during the Multiple Displacement Amplification reaction. BMC Biotechnol 7, 19 (2007).

31. Kiguchi, Y., Nishijima, S., Kumar, N., Hattori, M. & Suda, W. Long-read metagenomics of multiple displacement amplified DNA of low-biomass human gut phageomes by SACRA pre-processing chimeric reads. DNA Res 28 (2021).

32. Lu, N. et al. Exploration of whole genome amplification generated chimeric sequences in long-read sequencing data. Brief Bioinform 24 (2023).

33. Kermekchiev, M.B., Kirilova, L.I., Vail, E.E. & Barnes, W.M. Mutants of Taq DNA polymerase resistant to PCR inhibitors allow DNA amplification from whole blood and crude soil samples. Nucleic Acids Res 37, e40 (2009).

34. Sandberg, F., Kueng, N., Largiadèr, C.R. & Amstutz, U. Evaluation of variability in cell-free DNA extraction efficiency from plasma and urine and spike-in normalization. Sci Rep-Uk 15 (2025).

35. Maixner, F. et al. Linear polyacrylamide is highly efficient in precipitating and purifying environmental and ancient DNA. Methods Ecol Evol 13, 653–667 (2022).

36. Oreskovic, A., Brault, N.D., Panpradist, N., Lai, J.J. & Lutz, B.R. Analytical Comparison of Methods for Extraction of Short Cell-Free DNA from Urine. J Mol Diagn 21, 1067–1078 (2019).

37. Lundin, S., Stranneheim, H., Pettersson, E., Klevebring, D. & Lundeberg, J. Increased throughput by parallelization of library preparation for massive sequencing. PLoS One 5, e10029 (2010).

38. Xie, P. Model for forward polymerization and switching transition between polymerase and exonuclease sites by DNA polymerase molecular motors. Arch Biochem Biophys 457, 73–84 (2007).

39. Ganai, R.A., Bylund, G.O. & Johansson, E. Switching between polymerase and exonuclease sites in DNA polymerase epsilon. Nucleic Acids Res 43, 932–942 (2015).

40. Zhang, Z., Kermekchiev, M.B. & Barnes, W.M. Direct DNA amplification from crude clinical samples using a PCR enhancer cocktail and novel mutants of Taq. J Mol Diagn 12, 152–161 (2010).

41. McInerney, P., Adams, P. & Hadi, M.Z. Error Rate Comparison during Polymerase Chain Reaction by DNA Polymerase. Mol Biol Int 2014, 287430 (2014).

42. Deneke, J., Ziegelin, G., Lurz, R. & Lanka, E. The protelomerase of temperate Escherichia coli phage N15 has cleaving-joining activity. Proc Natl Acad Sci U S A 97, 7721–7726 (2000).

43. Gohl, D.M. et al. Systematic improvement of amplicon marker gene methods for increased accuracy in microbiome studies. Nat Biotechnol 34, 942–949 (2016).

44. Martinez-Garcia, E. et al. SEVA 4.0: an update of the Standard European Vector Architecture database for advanced analysis and programming of bacterial phenotypes. Nucleic Acids Res 51, D1558–D1567 (2023).

45. Calzadiaz-Ramirez, L. et al. In Vivo Selection for Formate Dehydrogenases with High Efficiency and Specificity toward NADP(). ACS Catal 10, 7512–7525 (2020).

46. Acinas, S.G. et al. Fine-scale phylogenetic architecture of a complex bacterial community. Nature 430, 551–554 (2004).

47. Baslan, T. et al. High resolution copy number inference in cancer using short-molecule nanopore sequencing. Nucleic Acids Res 49, e124 (2021).

48. Prabakar, R.K., Xu, L., Hicks, J. & Smith, A.D. SMURF-seq: efficient copy number profiling on long-read sequencers. Genome Biol 20, 134 (2019).

49. Di Tommaso, P. et al. Nextflow enables reproducible computational workflows. Nat Biotechnol 35, 316–319 (2017).

50. Cock, P.J. et al. Biopython: freely available Python tools for computational molecular biology and bioinformatics. Bioinformatics 25, 1422–1423 (2009).

51. Camacho, C., et al. BLAST+: architecture and applications. BMC Bioinformatics 10, 421 (2009).

52. Katoh, K. & Standley, D.M. MAFFT multiple sequence alignment software version 7: improvements in performance and usability. Mol Biol Evol 30, 772–780 (2013).

53. Ewing, B. & Green, P. Base-calling of automated sequencer traces using phred. II. Error probabilities. Genome Res 8, 186–194 (1998).

54. Maginn, E.J., et al. Monte Carlo methods, 70 years after “Equation of state calculations by fast computing machines” by Nicholas Metropolis, Arianna Rosenbluth, Marshall Rosenbluth, Augusta Teller, and Edward Teller (1953). J Chem Phys 163 (2025).

55. Kirkpatrick, S., Gelatt, C.D., Jr. & Vecchi, M.P. Optimization by simulated annealing. Science 220, 671–680 (1983).

56. Li, H. Minimap2: pairwise alignment for nucleotide sequences. Bioinformatics 34, 3094–3100 (2018).

57. Li, H. New strategies to improve minimap2 alignment accuracy. Bioinformatics 37, 4572–4574 (2021).

58. Katoh, K. & Frith, M.C. Adding unaligned sequences into an existing alignment using MAFFT and LAST. Bioinformatics 28, 3144–3146 (2012).

59. Goldfarb, T. et al. NCBI RefSeq: reference sequence standards through 25 years of curation and annotation. Nucleic Acids Res 53, D243–D257 (2025).

60. Schoch, C.L. et al. NCBI Taxonomy: a comprehensive update on curation, resources and tools. Database (Oxford) 2020 (2020).

